# AP-LASR: Automated Protein Libraries from Ancestral Sequence Reconstruction

**DOI:** 10.1101/2023.10.09.561537

**Authors:** James VanAntwerp, Mehrsa Mardikoraem, Nathaniel Pascual, Daniel Woldring

**Affiliations:** Department of Chemical Engineering and Materials Science, Michigan State University, East Lansing, MI 48824, USA; Institute for Quantitative Health Science and Engineering, Michigan State University, East Lansing, MI 48824, USA

## Abstract

**Background:** Ancestral sequence reconstruction (ASR) provides an informative roadmap of evolutionary protein sequence space that benefits protein design and engineering in pursuit <u>of high stability and diverse functionality</u>. Using statistical and biological knowledge, ASR can determine the most probable ancestor among potential alternative amino acid states. However, the inherent uncertainty of ASR can be further leveraged to determine viable “nearby” ancestors with wide-ranging functionalities by sampling alternative amino acid states.

**Results:** Here we introduce AP-LASR which i) automates ASR and ii) leverages uncertainty in ASR to generate diverse protein sequence libraries that consist of ancestral sequences and near-ancestor sequences. In addition to automating pre-processing tasks (e.g., data cleaning, multiple sequence alignment, and software dependency management), AP-LASR offers several user-definable hyperparameters (e.g., input data size, ancestral probability cut-off, and sequence supplementation) to control the properties of the generated library. AP-LASR features an improved eLnP score (a metric for quantifying reconstructed ancestral sequence confidence) compared to FireProt^ASR^, a well-established ASR workflow, for all four functionally diverse protein families studied. Furthermore, the rigorous statistical analysis undertaken in this study elucidates the influence of hyperparameters on ASR, enabling researchers to refine AP-LASR to their specific research.

**Conclusion:** AP-LASR offers an automated ASR experience that surpasses existing software by including a novel library design feature, powering curated protein libraries for wet-lab evaluation. We demonstrate how computational parameters impact the quality of ASR results, library composition, and the tradeoffs therein. AP-LASR offers a powerful tool for protein engineers to efficiently navigate the vast protein sequence landscape.

Software available at: https://github.com/WoldringLabMSU/AP-LASR

## Background

### 1.1. ASR is a powerful tool for protein engineering

The likelihood of random success is infinitesimal when designing proteins with improved or novel functions. The protein fitness landscape is rugged and sparse, with **the vast majority of protein sequences entirely lacking function**[1–4]. An efficient approach for navigating this space combines large collections of carefully chosen protein variants (i.e., libraries) with high-throughput experimental methods to rapidly characterize fitness. Choosing a robust starting point (i.e., a protein amenable to mutations) to explore the protein fitness landscape is an important consideration that is directly addressed by ancestral sequence reconstruction. Ancestral sequence reconstruction (ASR) is a method in bioinformatics that uses modern homologous sequences and statistical models of protein evolution to predict the common ancestors of modern proteins[5–7]. Importantly, ancestral proteins can be more thermally stable, more amenable to mutation, and more substrate-promiscuous than their modern homologs[7–15], making ancestral proteins attractive starting points for directed evolution and combinatorial library design

In protein engineering, ASR has been an instrumental technique in elucidating and advancing biological functions. A series of studies, including those by Thornton et al. and Bridgham et al., explored hormonal communication mechanisms, such as ancestral estrogen signaling and steroid hormone receptors evolution [7,8]Wheeler et al., Nguyen et al., and Akanuma et al. provided insights into thermostability, enzyme catalysis thermoadaption, and thermophilicity of ancestral life [9–11]. Protein G was effectively explored by Devamani et al., Starr and Thornton, Risso et al., and Harris et al., who investigated ancestral esterases, protein evolution epistasis, beta-lactamases, and cytochrome P450s, respectively [12–15]. Notably, Wilson et al. and Zakas et al. explored ASR in therapeutic areas, uncovering cancer drug mechanisms, and developing clinical coagulation factors[16,17]. In sum, these studies highlight ASR’s transformative influence on the protein engineering domain.

### 1.2. Uncertainty in ASR can be leveraged to identify viable regions of sequence space by introducing plausible amino acid diversity to most probable ancestors

As most of the sequences in the protein design fitness space lack function, the random mutations introduced through mutational library design generally produce destabilizing variants[1,2]. Traditional applications of ASR focus on identifying an individual protein sequence for each ancestor[7,11,12,15,18] but ASR can also generate a large collection (∼10^3^-10^9^) of plausible sequences for each ancestor. While a single amino acid can be inferred with high confidence at most positions of an ancestral protein, the uncertainty inherent to statistical predictions of the past can result in multiple amino acids being statistically plausible in key positions of the sequence. Thus, combinations of probable amino acids at multiple positions form a library of plausible ancestral sequences. Recently, Eick et al. [19]demonstrated that the stability and fitness of ancestral proteins can be maintained even when arbitrarily choosing less plausible amino acids at positions with multiple plausible amino acids. **This suggests that the regions of sequence space near the most probable ancestor are populated with diverse, functional proteins**. This feature of ASR provides an opportunity for designing large combinatorial libraries with the distinct advantage of maintaining sufficient stability and fitness among the library’s variants [18,20].

### 1.3. ASR is a non-trivial task

As we have described in a previous publication, ASR can be performed manually [20]. However, there are a number of barriers to performing ASR. Key tasks (e.g., sequence alignment, phylogenetic tree construction, and ancestor reconstruction) use separate software that generate output files that must be carefully processed and stored for subsequent steps. Furthermore, there are several nuanced considerations within each step that impact the overall performance of ASR. For example, in the initial step of curating a collection of modern sequences from public databases like UniProtKB, the dataset must be optimized to sufficiently cover a protein family by including relevant sequences while also minimizing computational demand of the workflow by excluding unnecessary sequences. In some cases, ASR may not be possible due to the lack of modern protein sequence data or the lack of computational resources. Automation lowers barriers to using ASR for library design.

### 1.4. ASR automation improves usability and accessibility. However, combinatorial library generation is not possible with current software

Given the challenges in manually performing ASR, various software (namely, MEGA11[21], FireProt^ASR^[22], and Topiary [23] have been developed in recent years to automate ASR. FireProt^ASR^ offers ASR through a convenient web server, returning a phylogenetic tree and ancestral data for a given sequence or multiple sequence alignment. In contrast, MEGA11 allows users to use local resources to perform more extensive analyses with a comprehensive suite of evolutionary biology tools and methods, including ASR. While Topiary can be used for ASR, it is primarily designed to create phylogenetic trees. Combinatorial libraries of ancestral sequences cannot be generated with any of these tools because of the limited access to underlying prediction confidences of inferred ancestors.

Here, we introduce AP-LASR, which automates an optimized ASR workflow, creates human-readable output metrics, and generates combinatorial protein libraries for wet-lab testing of the predicted proteins. Using consistent decision metrics, we performed ASR on four different protein families (i.e., estrogen receptor, dehalogenases, organic-anion transport peptides, all-trans-retinol dehydrogenases) under varying parameters to establish reasonable default user settings. In doing so, we demonstrated how qualities of the alignment (e.g., overall similarity, size, supplementation) affect the confidence distributions of the resulting predicted ancestral sequences. In comparison with the current best-in-class (FireProt^ASR^) method, AP-LASR features comparable or improved ASR quality while also generating diverse combinatorial libraries that enable efficient exploration of the rugged protein fitness landscape.

## Methods

In this section, we outline the steps taken by AP-LASR for automation of ASR and how it leverages uncertainty estimations in the reconstruction to create a protein library. We provide a justification for the choice of default parameters and explore the relationships between ASR initial parameters with the output quality metrics through a rigorous statistical analysis.

### 2.1 AP-LASR Workflow

When using AP-LASR, it is first necessary to download the Python script from GitHub (https://github.com/jjvanantwerp/Automated-ASR) and install the other software dependencies: MAFFT[24], CD-Hit[25], IQ-Tree[26], and the python modules Biopython [27] and xml.

### Ancestral Sequence Reconstruction

The AP-LASR workflow is initiated with the collection of related modern sequences based on a user-provided protein sequence or multiple sequence alignment for a selection of proteins. Using BlastP, AP-LASR identifies analogous sequences from NCBI’s non-redundant protein database. These sequences are then refined with CD-Hit to ensure diversity and compatibility with the input sequence length. Furthermore, the number of sequences included in this curation step may be further refined by a user-defined value. Subsequently, the sequences are aligned using MAFFT[24], and a phylogenetic tree is constructed with IQ-Tree[26]. This tree also infers ancestral sequences at various tree points, relying on a statistical model that traces the evolution of the protein family. A significant advantage of IQ-Tree is its ability to evaluate and select the optimal model of sequence evolution for the specific protein family under study[28].

If multiple sequences are provided as an input, AP-LASR will create a reconstruction showing all the connections between these input sequences. The specific sequences used in the input file may not be preserved in the final alignment, but sequences that cluster with them in CD-Hit will be. Finding where the input sequences fit into the final alignment simply involves reading the clustering information from CD-Hit (this is printed to the terminal as AP-LASR runs). This mode is especially useful if the common ancestor of two or more specific modern sequences is desired.

### Combinatorial Library Generation

Using the reconstructed ancestors and their uncertainties, combinatorial libraries can be generated. Figure 2A shows a phylogenetic tree with internal nodes representing predicted ancestral sequences. AP-LASR builds a library by first evaluating the statistical support for the topology of each internal node on the phylogenetic tree (i.e., tree support values). Of the ancestral nodes with good support values, libraries are generated using a confidence threshold approach. Each position within an ancestral sequence has a likelihood value for each of the twenty canonical amino acids. Typically, in ancestral sequences, most positions have a 99% or greater likelihood for either a gap or one amino acid, with a minority of positions having more uncertainty. Uncertain positions may have complicated predictions for amino acids, as shown in Figure 2B. From this distribution of probabilities, libraries are assembled using the threshold method: at each position, any amino acid with a percentage likelihood greater than the user-defined threshold is included in a combinatorial library. If no amino acid at this position meets the threshold, only the most likely amino acid is included. Higher thresholds, such as 20% likelihood, lead to much smaller libraries than lower thresholds, such as 5% likelihood. A summary of ASR analysis metrics and data files generated throughout the automated workflow are provided to the user as an output. For a representation of this workflow, see Figure 1.

**Figure 1.**
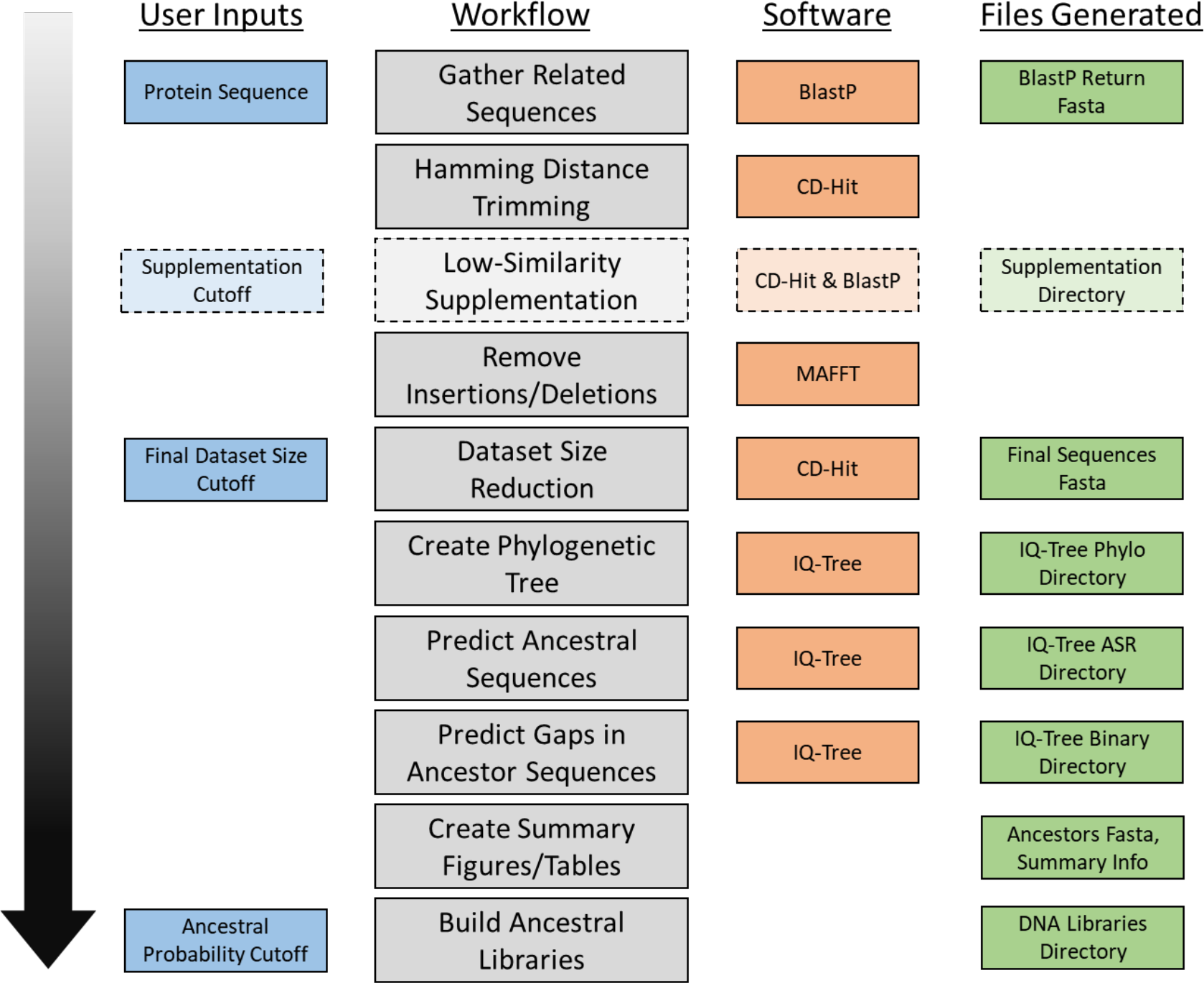
The workflow of AP-LASR, with tasks in the left column (grey), software used in the center-left colmn (salmon), user-defined variables in the center-right column (blue) and significant output file in the right column (green). Tasks are carried out in top-to-bottom order.

**Figure 2:**
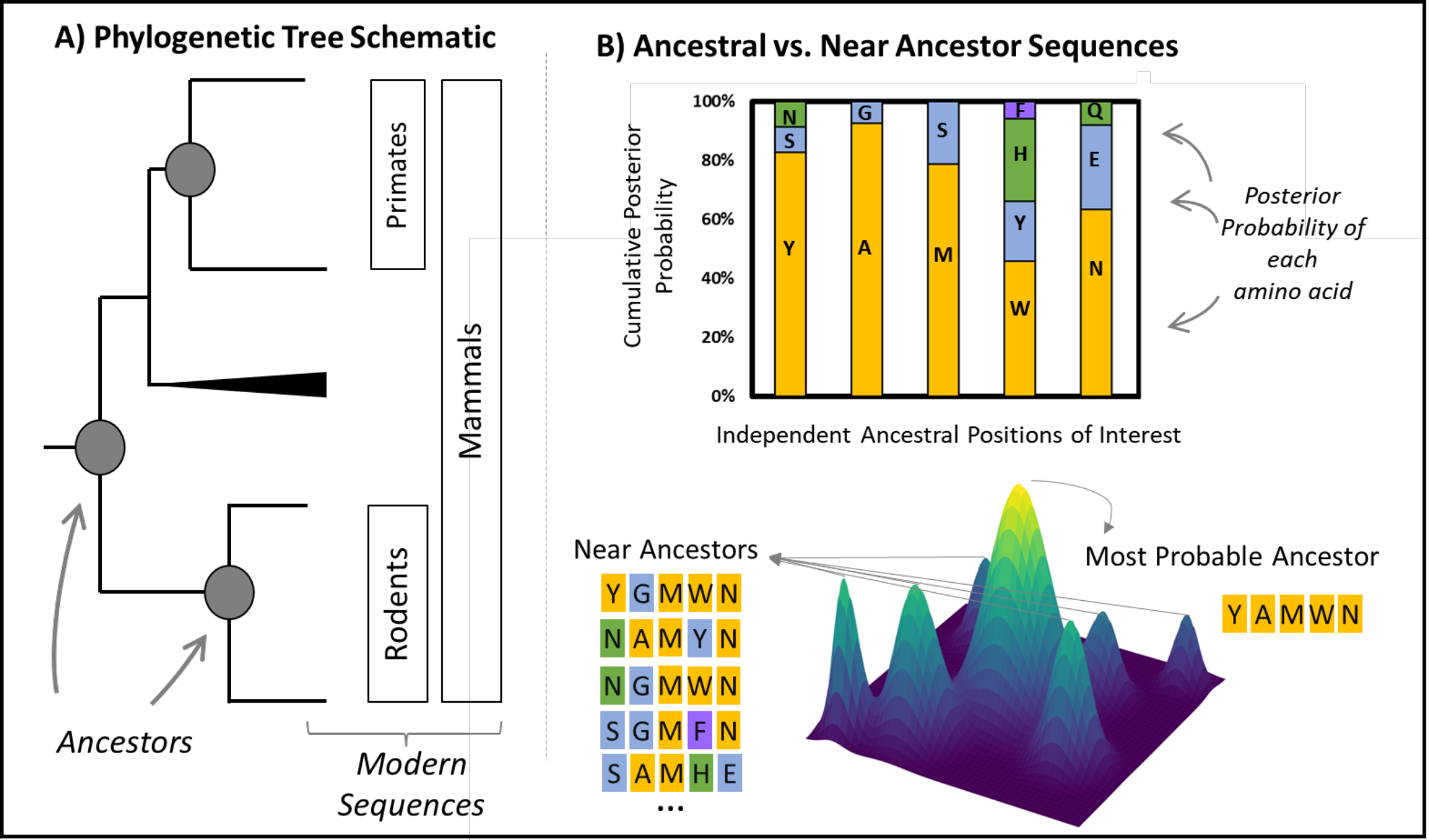
A) A phylogenetic tree, with predicted ancestral sequences at the internal nodes. B) The uncertainty in the prediction of an ancestral sequence. Each displayed letter represents the most likely ancestral amino acid, with the background color indicating confidence according to the legend. For ease of visualization, the lowest confidence sequence that could be found is displayed here.

### User-definable Parameters

As briefly described above, three user-defined parameters are included in AP-LASR to allow for flexibility to tailor ASR campaigns for users with different research objectives and various computational resource availability.

- **Ancestral probability cutoff:** Determines the confidence level in an ancestral residue for its inclusion in the combinatorial library. By default, AP-LASR produces eight libraries with varying cutoffs, but users can specify a particular cutoff.
- **Final MSA Dataset Size Cutoff:** Specifies the desired sequence count for phylogeny creation and ancestor reconstruction. Sequence clustering in CD-Hit reduces the dataset, ensuring no sequence similarity falls below 80%. The default is set at 500 sequences.
- **Supplementation Cutoff:** Defines the similarity level between a sequence and the dataset, qualifying the sequence for BlastP resubmission to enhance sequence coverage in distinct tree regions. This feature is typically OFF but can be activated based on specific requirements.

### 2.2 Experiment Setup & Decision-Support Methods

AP-LASR is designed to provide high-quality ASR with minimal user input; therefore, it is critical that the default parameters of AP-LASR permit high-quality ASR results for a diversity of proteins. We demonstrate here how input conditions affect the quality of final ASR, and what metrics can be evaluated to confirm results. The automation of ASR allowed testing of computational parameters in an unbiased way, examining how user-specified variables and the qualities of the underlying alignment support good ASR in general. It is also critical that we demonstrate that AP-LASR is comparable to or better than other software that already automate the ASR process. Here are some metrics we use for quantifying ASR quality and understanding the characteristics of the underlying alignment:

- Mean ASR Confidence: This is the average sequence confidence over all the most probable ancestor sequences in the reconstruction. Sequence confidence is the average confidence over each position in the sequence of the most likely amino acid state at that position. This metric shows the confidence in the reconstruction of ancestors, across the whole dataset.
- Mean SH-aLRT Support: This is a modification to the approximate likelihood-ratio test[29] measure of tree support first proposed by Shimodaira and Hasegawa[30].
- Mean UFB Support: This is a different measure of tree support that depends on random resampling of trees[31,32].
- Final Dataset Similarity: This percent similarity is a metric output by CD-Hit while clustering the sequences to increase the overall dataset diversity. At a given clustering cutoff, sequences are placed into a cluster where all sequences in the cluster are more similar than the cutoff. From each cluster, a representative sequence is chosen, and the reduced dataset will be only the representative sequences. The effect of this is that in the reduced dataset no two sequences are more than the cutoff percent similar to each other (if they were, they would be clustered together). This cutoff percentage is called the Final Dataset Similarity, so that a lower percentage is a more clustered dataset with fewer sequences than the unclustered dataset.

For AP-LASR parameter testing, we selected five proteins (refer to Table 1) due to their diverse sizes, functions, and structures, all of which are relevant for lab or clinical applications.

**Table 1:** Protein Families Under Study for the characterization of AP-LASR.

| Protein (Abbreviation) | UniProt ID | Protein Class | Organism of Origin | Available Homologs | Length (AAs) |
| --- | --- | --- | --- | --- | --- |
| Estrogen Receptor (ER) | Q99527-1 | Steroid Receptor GPCR | Human | Few | 375 |
| Dehalogenase (DH) | P59336-1 | Soluble Enzyme (Hydrolase) | Rhodococcus sp. | Many | 294 |
| Organic Anion Transport Protein 1B3 (OATP) | Q9NPD5-1 | Transmembrane Ion Transport | Human | Many | 702 |
| All-trans-retinol Dehydrogenase (ADH) | P08319 | Soluble Enzyme (Oxidoreductase) | Human | Many | 380 |

Two different input parameters were tested for each protein family: final allowed dataset size and supplementation cutoff. Preferred final dataset size was tested at 100, 200, 400, 800, and 1600 sequences. Supplementation was tested at supplementation similarities of 0% (no supplementation), 65%, 70%, 75%, 80%, and 85% (See the supplement for experimental conditions and results). <u>We used three metrics to assess the quality of the ASR outcome: ultra-fast bootstrapping (UFB) tree support, SH-aLRT tree support, and average ancestral prediction confidence</u>. However, using average values can be misleading since the distributions of these metrics often show a strong skew when considering all ancestors in a tree. Therefore, we base our conclusions on comparisons between these distributions and use statistical analysis to validate the significance of our findings.

### 2.3 Statistical Analysis

We implemented a rigorous statistical analysis in the experiment design runs between central parameters involved in the library design. This enables identifying potential correlations between user input parameters (e.g., dataset size, supplementation cutoff) and final ASR results such as mean ASR confidence, and mean UFB support. In addition, knowledge gained from this analysis benefits design fine-tuning in specific ASR use cases. To identify the statistically significant correlations among the library design parameters, a permutation test was applied. The permutation test is a non-parametric statistical test and was used to test the null hypothesis that there is no significant correlation between the two chosen variables (i.e., AP-LASR experiment design parameters). This non-parametric test enables identifying correlations without making assumptions about the underlying distribution of the dataset and is viable in low sample sizes.

### 2.4 Expected Log-probability (eLnP) to benchmark ancestor reconstruction reliability

The reliability of the final reconstructed ancestry is quantified using the eLnP metric, as proposed by Sennett and Theobald (Equation 1)[33]. eLnP quantifies the overall uncertainty in the reconstructed sequence distribution and was used as our benchmark to evaluate the efficacy of AP-LASR against well-established software such as FireProt^ASR^.

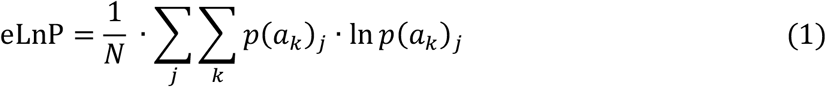

In Equation 1, *a* symbolizes an amino acid state, *k* is one of the twenty amino acids, and *j* designates its position within the sequence of length *N*. By aggregating probabilities across amino acids and positions, we derive a thorough evaluation of sequence confidence.

## 3 Results and Discussion

### 3.1 AP-LASR Achieved Higher Confidence Score Compared to FireProt^ASR^

Across all protein families tested, AP-LASR achieved improved or comparable eLnP scores compared to FireProt^ASR^. Note that a greater (less negative) eLnP number represents greater confidence[33]. This metric was chosen for two reasons: it can compare ASR outputs from various software or models in an unbiased way and it offers assessments even in the absence of definitive ‘correct’ ancestors. Figure 3 distinctly showcases the eLnP distribution for sequences obtained via AP-LASR in comparison to those from FireProt^ASR^. The higher eLnP values, and theie reduced standard deviation, for AP-LASR indicate an elevated confidence in its reconstructions. This analysis demonstrates that AP-LASR performance in reconstruction is comparable and in some cases was improved compared to FireProt^ASR^, well-known for its reliability and precision for ASR.

**Figure 3:**
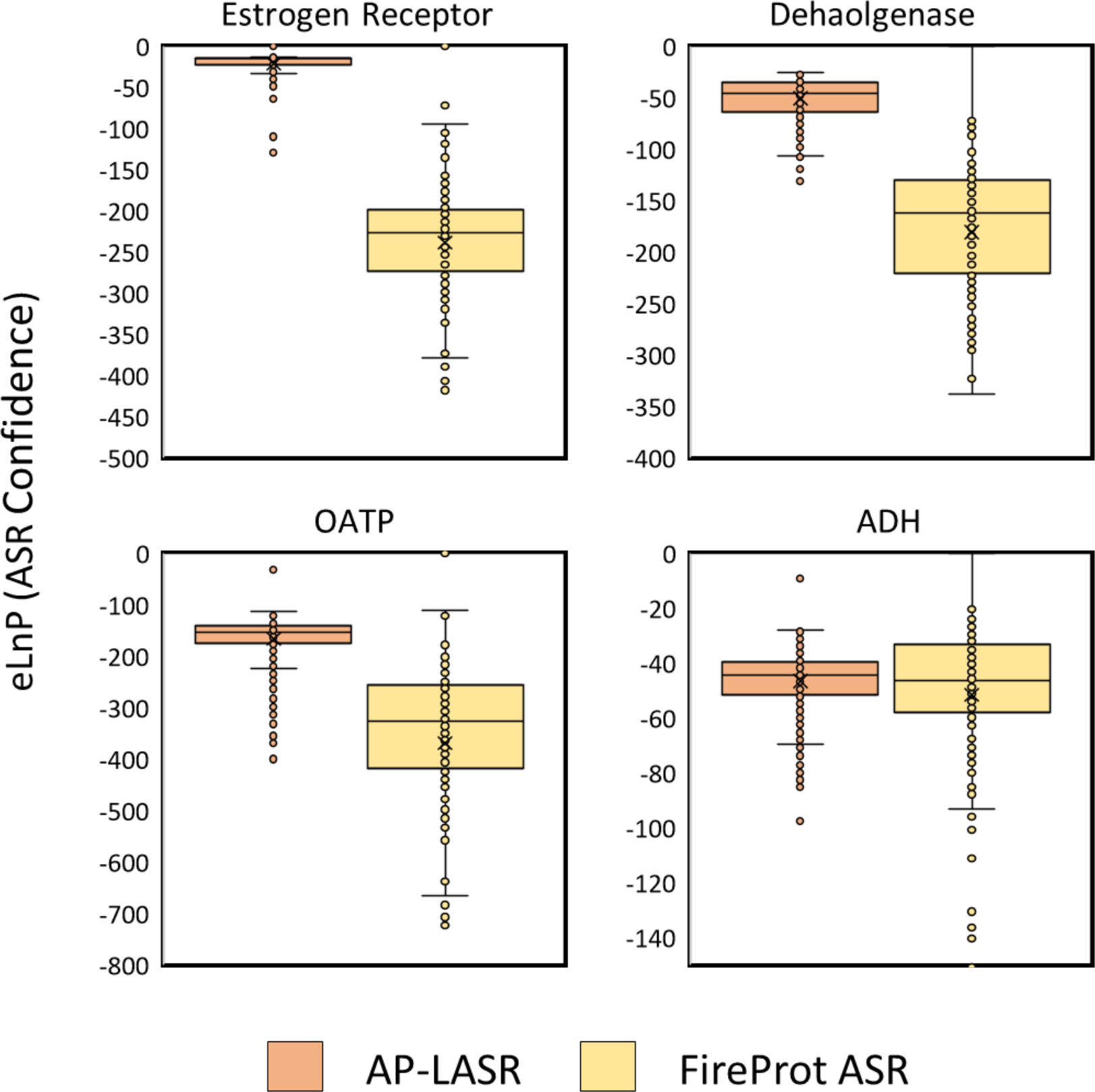
eLnP score comparisons obtained via reconstruction of each protein family which was consistently higher in AP-LASR. Note that AP-LASR reconstructions contained no more than 400 modern sequences.

### 3.2 Library Size and Ancestral Probability Cutoff

The length of the ancestral sequence, uncertainty in the prediction of a particular ancestral sequence, and ancestral probability cutoff all determine the size of the resulting library. Figure 4 shows how ancestral probability cutoff affects the library size for different ancestors. All libraries grow factorially because of their combinatorial nature, but the connection between confidence threshold and library size depends on the inherent uncertainty of the predicted ancestral sequence. Looking at the reconstructed OATPs in Figure 4 Node 322 never creates a library larger than 12,300 sequences because of its high prediction confidence. The library of Node 189 grows to astronomical size because of the significant uncertainty in prediction of that sequence.

**Figure 4:**
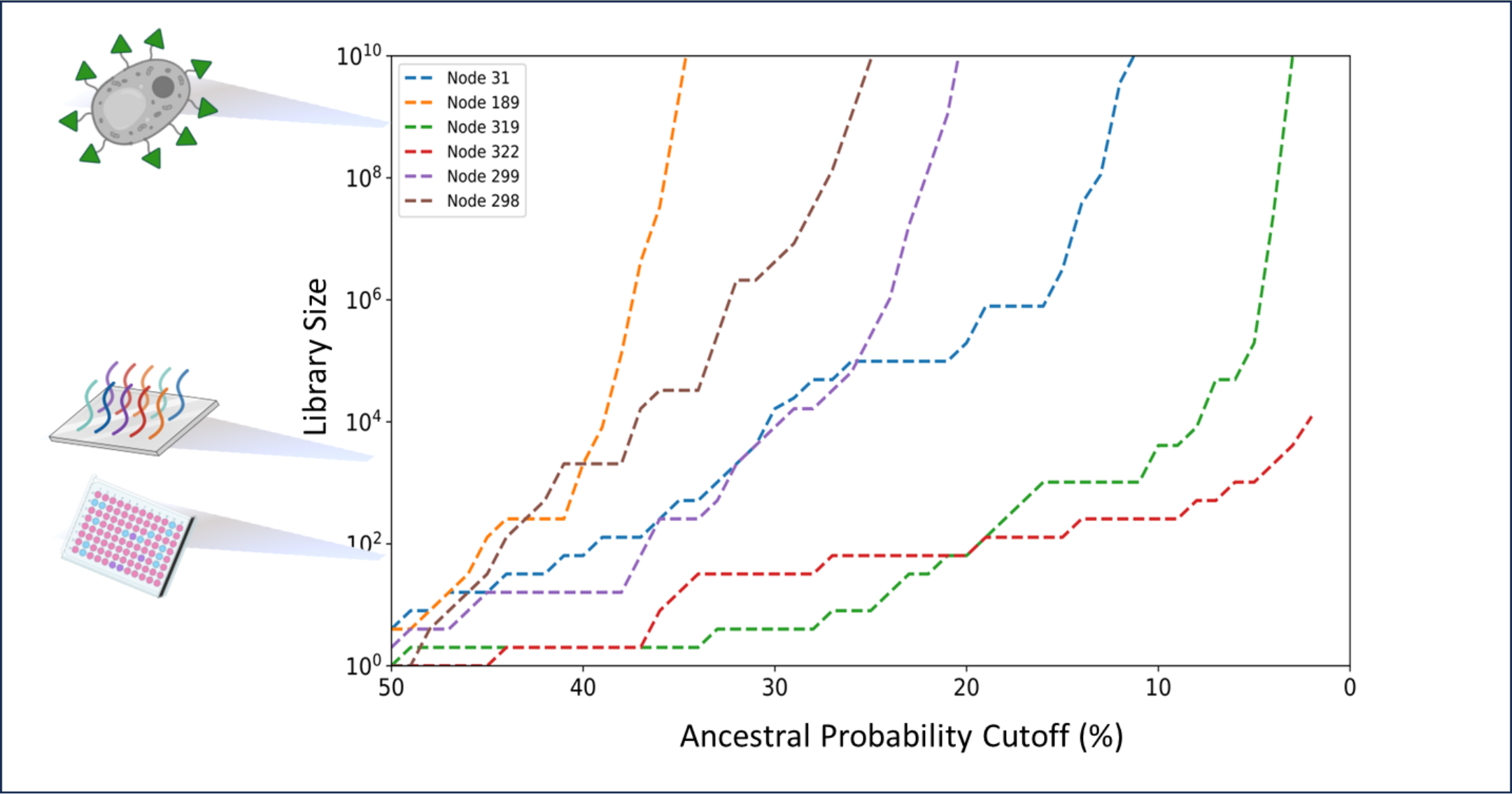
Depiction of selected ancestral nodes derived from the OATP reconstruction tree, illustrating the influence of library size and confidence threshold. The confidence threshold can be optimized based on the experimental throughput (e.g., plate reader (E2), microarray (E3), yeast display (E9), phage display (E12)).

For every sequence, when the ancestral probability cutoff is reduced, more mutations are added to the combinatorial library and as a result the number of near ancestor sequences substantially increases. This effect is amplified by the skewed distribution of ancestral states: there are more alternative states with low cutoff/confidence than with high cutoff/confidence. As a result, even minor adjustments to the probability cutoff, especially at the lower end, can dramatically affect the library’s final size. This trend was shown in all the protein families tested in this study. This parameter can be optimized based on the protein family and experimental approach to evaluating the functionality of candidates in the final library. By default, AP-LASR will generate a range of libraries at a range of cutoffs and list the size of each library for each ancestor. Libraries can be re-generated at a specified cutoff using the “RemakeLibraries” mode.

### 3.3 Influence of Sequence Similarity and Dataset Size on Ancestral Sequence Reconstruction Confidence

CD-Hit is implemented in AP-LASR to group sequences into distinct clusters, with only a representative sequence from each cluster being retained for the final dataset (as illustrated in Figure 5A). As the clustering threshold increases, there’s a noticeable increase in the dataset size curated for ancestral reconstruction. Figure 5B showcases the effect of varying the CD-Hit similarity threshold across our four distinct protein families. While an undeniable increase in final dataset sizes is observed, 0’s significant to note that different protein families have different amounts of data available, affecting the domain of the curve.

**Figure 5:**
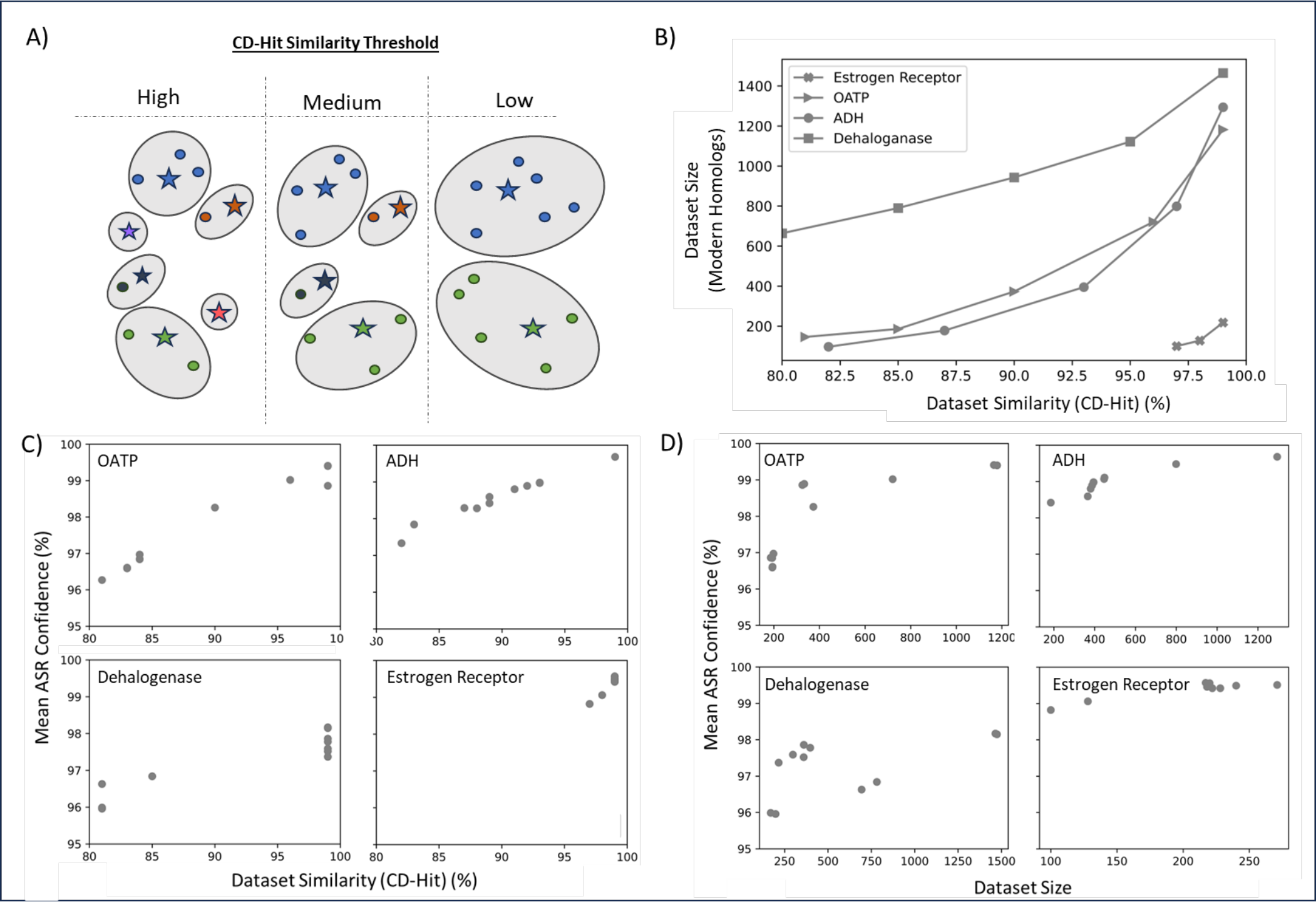
Interplay of dataset similarity cutoff and dataset size on the fidelity of reconstructed sequences in AP-LASR. A-Schematic representation of the effect of increasing CD-Hit threshold on the final sequences. This threshold dictates the required similarity of sequences to be clustered in the same group. The higher the threshold, the less probable the sequences clustered in the same group resulting in less final diversity and a greater number of sequences. B-Increased CD-Hit results in rising dataset size yet different protein families have different growth rates. For instance, the dehalogenase family reaches a point where sequences become too different to cluster further, whereas the estrogen receptor family has a limit of approximately 220 sequences based on available BLAST-P results. C-Mean ASR confidence rises with increased similarity among input sequences. D-Comparison of ASR confidence improvements with dataset size across various protein families. Notably, the enhancements are not consistently significant across all families. For the OATP protein family, a larger dataset size leads to better average ASR confidence. This specific case highlights the significant boost in ASR confidence with increased dataset size. It’s worth noting that AP-LASR’s default setting is 500 sequences in the alignment and has shown to be sufficient in our experiments for reliable ASR. While adding more sequences can enhance ASR, the benefits tend to taper off with much larger datasets.

Our statistical analysis underpins that the correlation between CD-Hit clustering and ASR confidence is significant. This suggests the chosen CD-Hit similarity threshold plays a crucial role in determining the reliability of ancestral sequence predictions (i.e., ASR confidence). On the other hand, while the dataset size is inherently influential, its correlation with ASR confidence isn’t universally significant. However, for specific protein families, such as OATPs, a significant correlation exists. This is likely because of the tie between dataset size and similarity cutoff (Figure 5B). With a more similar dataset, there are fewer mutations to infer between adjacent proteins in the phylogenetic tree, improving confidence in ancestral sequences. Interestingly, these is a weak inverse correlation for tree supports; section 3.4. Given these insights, 0’s paramount to curate a dataset that’s not simply large but rather representative, diverse, and reasonably dense. The interplay between dataset size, ASR confidence, and similarity thresholds is shown in Figures 5C and 5D. All correlations and their statistical confidence can be found in the supplement.

### 3.4 Tree Support Metrics are Positively Correlated with Dataset Diversity and Negatively Correlated with Measures of Alignment Distance

Tree support values, UFB and SH-ALRT which indicate the robustness of phylogenetic tree, reveal a positive correlation with dataset diversity attributes measured by hamming distance distribution, highlighting their reliance on the underlying data characteristics. Both UFB and SH-aLRT values are associated with the mean Hamming distance of a dataset (ρ=0.89 and ρ=0.80, respectively), as well as the median Hamming distance (ρ=0.61 and ρ=0.69, respectively). There is also a positive correlation between the percentage of alignment gaps and the tree support values (ρ=0.65 and ρ=0.58). The findings suggest that an increased number of gaps and a greater Hamming distance leads to phylogenetic trees with better support values, possibly due to reduced ambiguity in tree assembly under such conditions; the placement of more similar sequences on the tree is more ambiguous. In this way, there is a tradeoff between tree confidence and ASR confidence.

### 3.5 Balancing ASR confidence & Tree Support Values

In IQ-TREE analysis, the relationship between Ancestral Sequence Reconstruction (ASR) confidence and tree support metrics, UFB and SH-aLRT, was investigated. Initially, Spearman correlation coefficients of -0.39 and -0.50 were observed, suggesting a moderate negative correlation. However, permutation tests results were not statistically significant, implying that the observed relationships might have occurred by chance. A practical approach is to build a robust phylogenetic tree and obtain best supported branches is using cutoff SH-aLRT > 80% and UFB > 95%, (recommended by IQ-Tree) while also aiming for a high ASR confidence. Aiming to maximize ASR confidence results in fewer near-ancestor sequences in the final pool for experiments. Therefore, depending on the wet-lab throughput and the capacity to handle uncertainty by incorporating more near ancestors, users may need to fine-tune the confidence threshold for the final library.

## Conclusions

Here we present AP-LASR, a software for the automated generation of protein libraries from automated ancestral sequence reconstruction. We hope that this software will make the process of ASR faster and easier and will allow many more protein engineers to build libraries tailored to the mutational space of their protein and their expression method. To this end, we tested the impact of AP-LASR’s parameters on measures of ASR results and determined that there is a trade-off between ASR confidence and tree support values. We also determined that the supplementation feature which we expected to be helpful did not have a significant impact on the resulting ASR. We demonstrate that AP-LASR is comparable to the best-in-class automated ASR software.

## Supporting information

Experimental Setup and Results (xlsx)

## Availability of Data and Materials

Project Name: Automated ASR

Project Home Page: https://github.com/WoldringLabMSU/AP-LASR

Operating System: Linux or MacOS

Programming Language: Python

Other Requirements: IQ-Tree 2, CD-Hit, MAFFT.

License: MIT License (posted on GitHub)

## Notes

https://github.com/WoldringLabMSU/AP-LASR

### Competing Interest Statement

The authors have declared no competing interest.

